# KBeagle: An Adaptive Strategy and Tool for Improvement of Imputation Accuracy and Computing Efficiency

**DOI:** 10.1101/2022.10.22.513369

**Authors:** Jie Qin, Xinrui Liu, Yaxin Liu, Wei Peng, Yixi Kangzhu, Jincheng Zhong, Jiabo Wang

## Abstract

With the development of molecular biology and genetics, deep sequencing technology has become the main way to discover genetic variation and reveal the molecular structure of genome. Due to the complexity of the whole genome segment structure, a large number of missing genotypes have appeared after sequencing, and these missing genotypes can be imputed by genotype imputation method. With the in-depth study of genotype imputation methods, computational intensive and computationally efficient imputation software come into being. Beagle software, as an efficient imputation software, is widely used because of its advantages of low memory consumption, fast running speed and relatively high imputation accuracy. K-Means clustering can divide individuals with similar population structure into a class, so that individuals in the same class can share longer haplotype fragments. Therefore, combining K-Means clustering algorithm with Beagle software can improve the interpolation accuracy. The Beagle and KBeagle method was used to compare the imputation efficiency. The KBeagle method presents a higher imputation matching rate and a shorter computing time. In the genome selection and heritability estimated section, the genotype dataset after imputed, unimputed, and with real genotype show similar prediction accuracy. However the estimated heritability using genotype dataset after imputed is closer to the estimation by the dataset with real genotype. We generated a compounds and efficient imputation method, which presents valuable resource for improvement of imputation accuracy and computing time. We envisage the application of KBeagle will be focus on the livestock sequencing study under strong genetic structure.

## Introduction

Following the development of molecular biology and genetics, the deep sequencing technology has been the major approach to discovery heritable variation and reveal molecular structure of genome. Based on the complicated construction of fragments in the whole genome, such as long replications, instability single-stranded structure, and the mistakes of duplication, there have been lots of missing genotype after sequencing. The drawback of having low density markers or more missing genotype for a sequencing study can be overcome by applying imputation. The principle of genotype imputation is to use the known genotype information in the reference panel to infer the possible genotypes in the target sample, usually using the linkage information near the missing genotype (Chen and Shi 2019). There are several kinds of genotype imputation software, which can be divided into computationally intensive or computationally efficient, depending on the length of imputing time. Computational-intensive imputation software uses all known genotypes to infer missing genotypes during imputation, thus, it takes longer computing time but has higher imputation accuracy. Such software includes IMPUTE2 (Howie et al. 2009), MaCH (Li 2006), fastPHASE (Scheet and Stephens 2006).

Computing efficient imputation software uses only the subset of genotypes adjacent to specific single nucleotide polymorphism (SNP) sites in the imputation process, resulting in lower imputation accuracy but shorter computing time. Such software includes PLINK (Purcell et al. 2007) and Beagle (Browning and Browning 2007).

As one of the imputation software, Beagle has three advantages over other imputation software. The genotype file input by Beagle software is in bref3 format, and the number of gigabytes required for reference sample data of the same size stored in bref3 format is smaller than other formats (Browning et al. 2018). Therefore, one of the advantages of Beagle software is its low memory consumption. Another advantage of Beagle software is that it runs faster. When using the same reference panel size, the Beagle software costs less times than the impute software and Minmac software (Browning et al. 2018). In addition, the imputation accuracy of Beagle is relatively higher. Because of the advantages of Beagle software, it is widely used in the study of genotype imputation of animals such as humans (Rubinacci et al. 2020), cattle (Chud et al. 2015; Uemoto et al. 2015), pigs (Song et al. 2019), chickens (Ye et al. 2019).

The purpose of clustering is to divide a dataset into multiple clusters (or classes) with high similarity within clusters and low similarity between clusters (Liu et al. 2021). In population genetic analysis, clustering is used to divide groups with a large number of individuals into groups with similar population structures. Stachowicz et al. (2013) reported that the stronger the correlation between the reference panel and the target sample, the higher the imputation accuracy. Clustering classifies strongly correlated samples into one category, resulting in longer haplotype fragments shared by highly correlated samples, and more accurate imputation results. Therefore, the combination of K-Means clustering algorithm and Beagle software may improve the imputation accuracy.

In this study, we aimed to compare the imputation matching rate and computing time of Beagle software and the KBeagle method. Next, we performed genome selection for the data imputed by the KBeagle method and calculated heritability and prediction accuracy to explore the imputation results of the Kbeagle method. The successful application of KBeagle method provides a new direction for the research of genotype imputation method in the future.

## Materials and Methods

### Genotype Data

This study referred to a previous study that included 354 Ashdan female yaks and 98K SNPs. The data (Jia et al. 2020) link is (https://www.animalgenome.org/repository/pub/NWAU2019.0430/). Three types of files were obtained: ‘ped’ file, ‘map’ file, and phenotype file. The ‘ped’ file contained 98688 SNPs, the ‘map’ file contained genetic map information, and the phenotype file contained four growth traits, all obtained from 354 yaks. The PLINK 1.9 software (http://www.cog-genomics.org/plink2) was used to convert ‘ped’ files into numeric format genome information files. The genotypes were coded as 0, 1, and 2, corresponding to major alleles, heterozygous genotype, and minor alleles, respectively.

Based on the original dataset, we divided the whole data into several sub-datasets: 10K, 30K, 50K, and 70K. They were randomly selected from 98K data, with each data size generating 30 replications. Missing genotypes were randomly sampled in proportions of 5%, 10%, 15%, and 20% and replicated 30 times. The missing genotypes were represented by ‘NA’, and the data containing the missing genotypes were used as the reference dataset. The missing genotypes in the reference dataset were imputed using the Beagle method and the KBeagle method. The missed genotype data was saved in another independent document as the true genotype of the prediction dataset.

## Imputation Algorithm

### Beagle

Beagle 5.1 used hidden Markov models (HMM) for genotype imputation. In the Li and Stephens (2003) model, the HMM state space is a matrix of reference alleles, with each row recorded as a reference haplotype, each column recorded as a marker, and each HMM state labeled with the allele carried by the reference haplotype at the marker. For each marker, the sum of the probabilities of the states labeled by an allele was the imputed probability for that allele (Browning et al. 2018). The probability of a specific haplotype was calculated by adding the probabilities from the first marker to the last marker. The haplotype set was obtained by genotyping the target samples. The model was established according to the common gene sequences between the target sample and the reference panel. The missing alleles in the target sample were predicted and imputed using the calculated probability of allele markers in the reference panel.

### KBeagle

The principle of the KBeagle method is to use the K-Means algorithm to calculate the genetic distance of samples with missing genotypes, classifying the samples with close genetic distances into one clustered group, and then use the Beagle software to estimate the missing genotype of samples in each clustered group. The K-Means algorithm divided the population into K cluster groups, each cluster group with several uncertain samples. This study assumed that the population was divided into 10 cluster groups, and the function fviz_nbclust in the factoextra package was used to find the best cluster number K less than or equal to 10.

The whole KBeagle progress consisted of 6 steps: Data Inputting, K-Means Clustering, Clustering Result, Beagle Imputing, Imputing Result, and Data Outputting (Fig 1). Numeric data formats could be accepted as input data. The whole data should include the taxa and the genotype with the missing value. The genotypes such as homozygotes of major and minor alleles, and heterozygotes should be coded as 0, 2, and 1 respectively (Fig 1A). Genome wide markers were used to estimate the genetic distance between each sample. Additionally, the K-Means algorithm was used to select K cluster centers, calculate the distance between the cluster center and the samples, and group those closest to the cluster center into one class. All samples were clustered into K classes (Fig. 1B). Next, all samples were reordered according to the clustering results, samples in the same cluster together were arranged, and K data sets were generated for subsequent imputation (Fig. 1C). The Beagle software was used to impute the samples in each data set (Figure 1D). K data sets containing all genotypes were imputed with Beagle software (Fig. 1E). The input samples were arranged in the original order and outputted according to the ID order of the input data (Fig. 1F).

**Figure 1.**
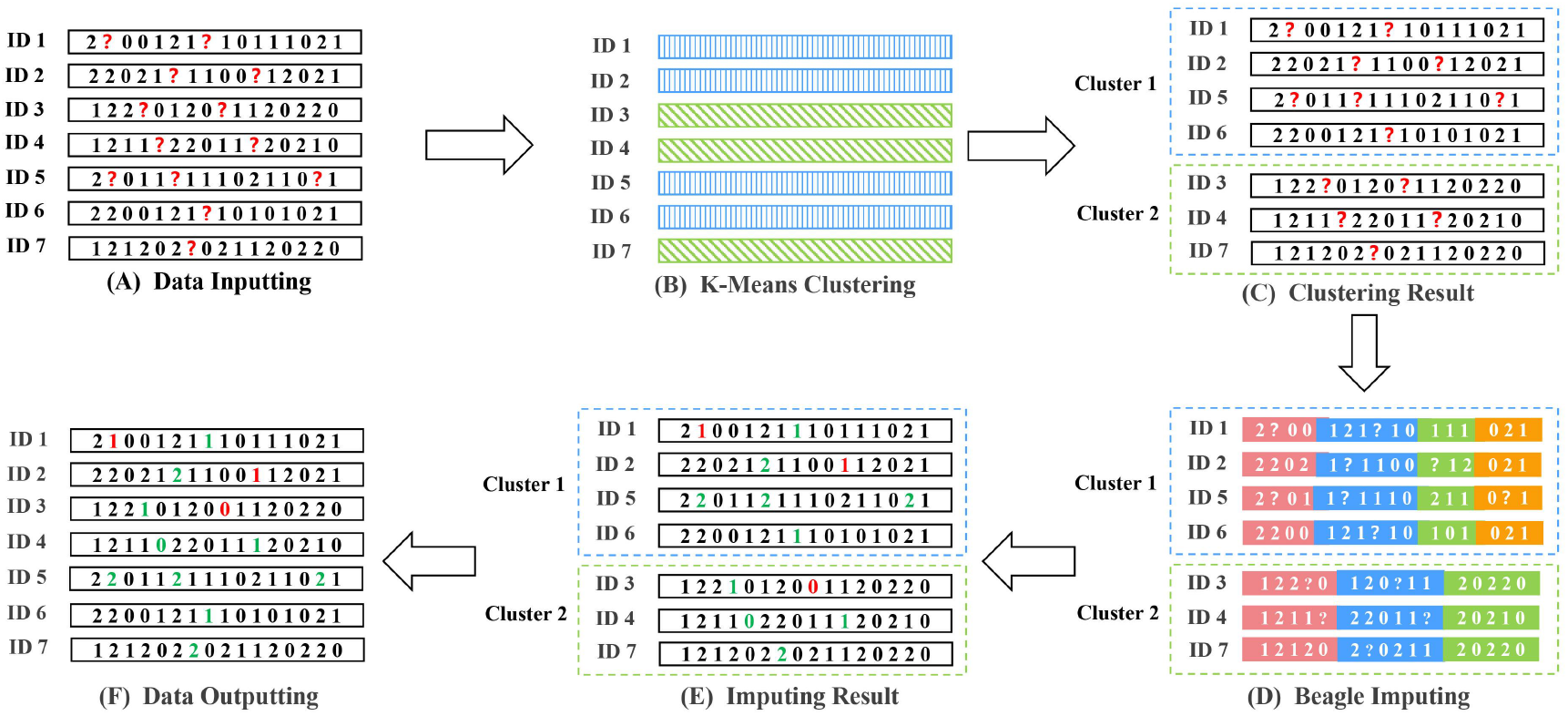
The pipline of data flow in the K-Means Beagle (KBeagle). In the KBeagle, the whole individuals with missing genotype were estimated genetics distance at first. Based on the genetics distance the whole population were divided into several clustered groups. The K-mean algorithm was used to cluster. Then the genotype of individuals in each clustered group were imputed using Beagle algorithm. Finally, the results after imputation will be ordered as previous. The whole KBeagle progress contained 6 steps: (A)Data Inputting. The numeric data format could be accepted as input data. Whole data should include the ID and genotype of each sample. The genotype such as homozygotes of major and minor alleles, and heterozygous should be coded as 0, 2, and 1 respectively. (B) K-Means Clustering. Whole genome markers were used to estimated genetics distance between each sample. Using K-mean algorithm individuals were clustered into groups. (C) Clustering Result. All samples were reordered according to the clustering results, the samples were arranged in the same cluster together, and generated K sub datasets (K was the number of sub datasets). (D) Beagle Imputing. Beagle software was used to impute the samples in each set respectively. (E)Imputing Result. K datasets containing complete genotypes were imputed. (F)Data Outputting. According to the ID order of the input data, the imputed samples were arranged in the original order and output.

### Definition of imputation ability

To compare the imputation ability of the two genotype imputation methods, this study used the imputation matching rate and computing time as the evaluation criteria. The imputation matching rate was referred to as the ratio of the number of correctly imputed genotypes to the number of totally imputed genotypes. The calculated formula is

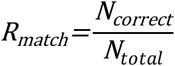

where *N*_*correct*_ was the total number of genotypes accurately predicted after imputation, and *N*_*total*_ was the total number of all missing genotypes.

Computing time was defined as the time it took from the beginning of imputation to the end of imputation, including data inputting, data processing, genotype imputation and the calculation of imputation accuracy. The system.time function started measuring the time from the data inputting step and ended when the imputation accuracy was calculated, thus, the total time of all steps was the computing time.

## Genome selection

### Simulated phenotype

In this study the GAPIT (Wang and Zhang 2021) (www.zzlab.net/GAPIT) software package Gapit.Phenotype.Simulation was used to simulate the population phenotype. In order to evaluate the effectiveness of genome selection using the imputed genotype data, the phenotypic data was simulated using the genotype data with missing values. Before running the simulation, we needed to calculate the missing rate of each SNP and arrange the misssing rate from high to low. When simulating the phenotype, the heritability was set to 75%, the number of quantitative trait nucleotides (QTNs) was set to 20, and the loci of QTNs were selected from the SNPs with the highest missing rates. The simulated phenotypic and genotypic data were then subjected to genomic selection analysis.

### Prediction model

The full name of rrBLUP (Medina et al. 2021) is ‘ridge regression best linear unbiased prediction’, which is the best linear unbiased prediction of ridge regression. It is one of the most commonly used models for genome selection and also represents the indirect method models. The principle of the indirect method is to estimate the marker effect in the reference population, and then combine the genotype information of the predicted population to accumulate the marker effect and to obtain the individual estimated breeding value of the predicted population.

The rrBLUP package mainly included two functions: the A.mat function and the mixed.solve function. The A.mat function was used to filter and impute genotypes, and the mixed.solve function directly solved model parameters. The mixed.solve function was the core of the rrBLUP package, and the solution (Endelman 2011) form was

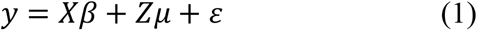

where *y* was the phenotype vector, *x* was the coefficient matrix of the fixed effect, β was the estimated fixed effect, *z* was the coefficient matrix of the random effect, *μ* was the estimated random effect, *ε* was the residual error, and followed the distribution 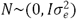.

### Estimated Heritability

In this study, narrow sense heritability (*h*^*2*^) was defined as the rate of additive variance (*V*_*A*_) in the total phenotypic variance (*V*_*p*_), and the (Wang et al. 2017) formula used was

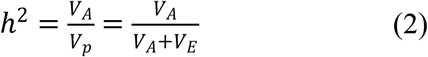

First, the phenotype was simulated using the Gapit.Phenotype.Simulation function in the GAPIT software package. Then, the real data, the data containing the missing genotypes and the imputed data were analyzed by genome selection using the GAPIT software package, and the heritability of the three datasets was calculated, and selected the gBLUP model for calculation. Each dataset was processed 30 times, and the average of the 30 replicates was taken as the heritability of each data.

### Prediction accuracy

Prediction accuracy was defined as the Pearson correlation coefficient between the predicted phenotypic value and the true phenotypic value with higher values indicating higher accuracy. First, the simulated phenotype data should be randomly divided into training and test phenotypes, and the corresponding genotype data should be divided into training and test sets, according to the classification results. The model was then built in the mixed.solve function of the rrBLUP package in R, substituting the training phenotype and training set, and selecting the maximum likelihood method for the calculation. Then, the test set was multiplied by the coefficient *μ* in the model to obtain the predicted phenotype value. Therefore, the Pearson correlation coefficient between the predicted phenotype value and the test phenotype gave the prediction accuracy. This was repeated 50 times and the average value was used to obtain the average accuracy of cross validation. Finally, all the data were repeated 30 times, and the average value was obtained to determine the average accuracy of all the data.

## Results

### Imputation matching rate

The imputation matching rate between the real dataset and the predicted dataset was calculated, each data set was repeated 30 times, and the average value was used as the final imputation matching rate of each method (Table 1, Fig 2). At 10K (Fig. 2A), 30K (Fig. 2B), 50K (Fig. 2C), and 70K (Fig. 2D), the imputation matching rate of the KBeagle method was always higher than that of the Beagle method. The imputation matching rate of the KBeagle and the Beagle methods decreased as the missing rate increased. Among the results of 10K, 30K, 50K, and 70K data, the imputation matching rate of data with 5% missing rate was the highest, whilst data with a missing rate of 20% had the lowest imputation matching rate. As the data set increased, the imputation matching rate also increased, with the imputation matching rate of 10K data being the smallest and the imputation matching rate of 70K data being the largest.

**Figure 2.**
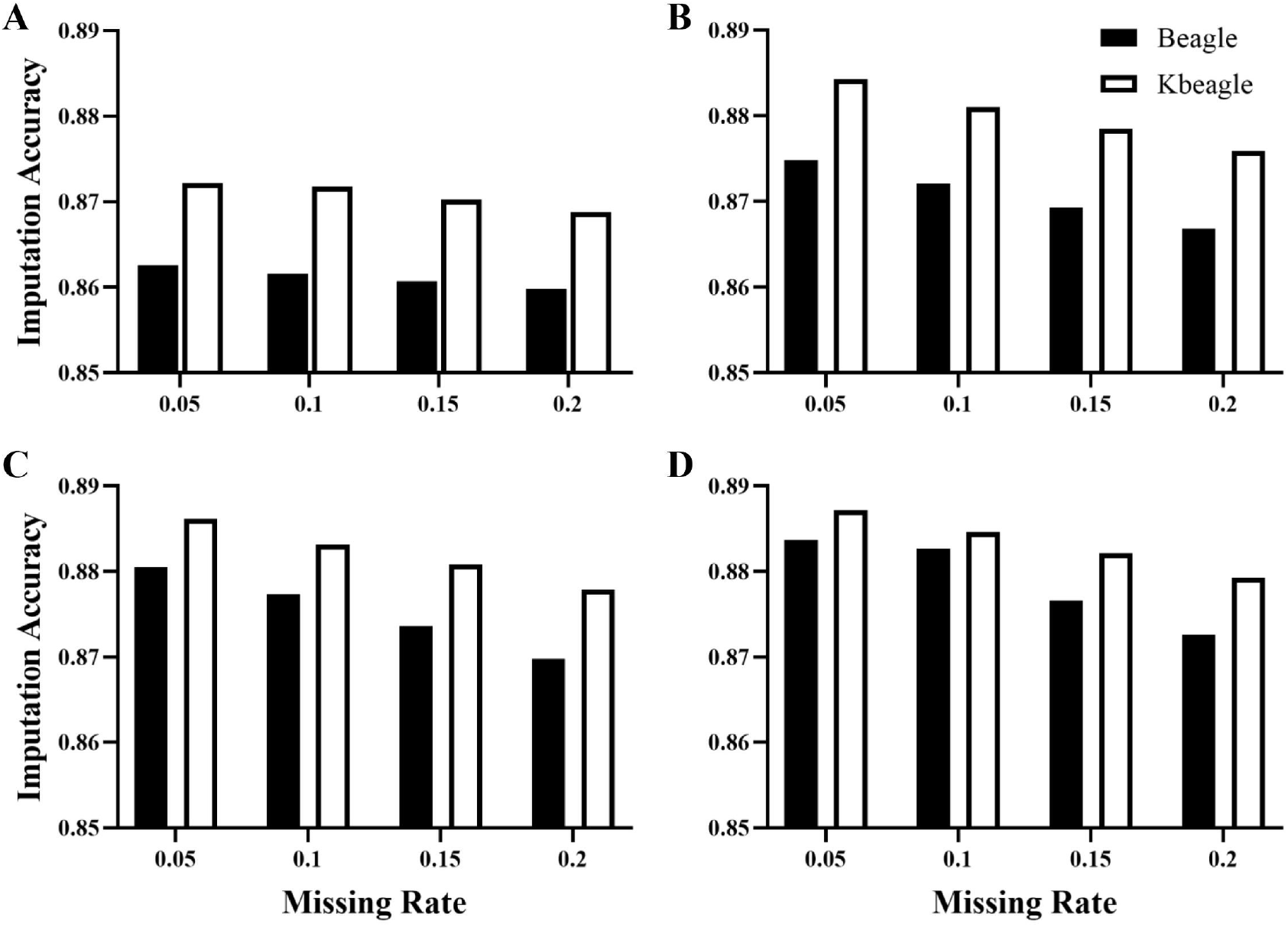
Assess of imputation accuracy between Beagle and KBeagle method. 10K(A), 30K(B), 50K(C) and 70K(D) data were randomly selected from the 98K data of 354 yaks, and each size of data generated 30 times replication of real data for genotype imputation. The data of each group were artificially missed genotype according to the proportion of 5%, 10%, 15% and 20% as reference data set. And the data containing the missing genotype was used as the real inference data set. The reference dataset was used to estimate parameters of Beagle and KBeagle model. Then based on these parameters the genotype in the inference were imputed. The matching ratio of the number of correct genotypes in the total number of missing genotypes was calculated as the imputation accuracy. The y-axis in each figure represented the imputation accuracy, the x-axis represented the missing rate, and the Beagle (blue bar) and KBeagle (red bar) were displayed as the average of 30 random replicates of data.

### Computing time

When using the Beagle and KBeagle methods for genotype imputation, time function in the R package was used to record the computing time of each time, and the average value was taken as the final computing time (Table 2, Fig 3).

**Figure 3.**
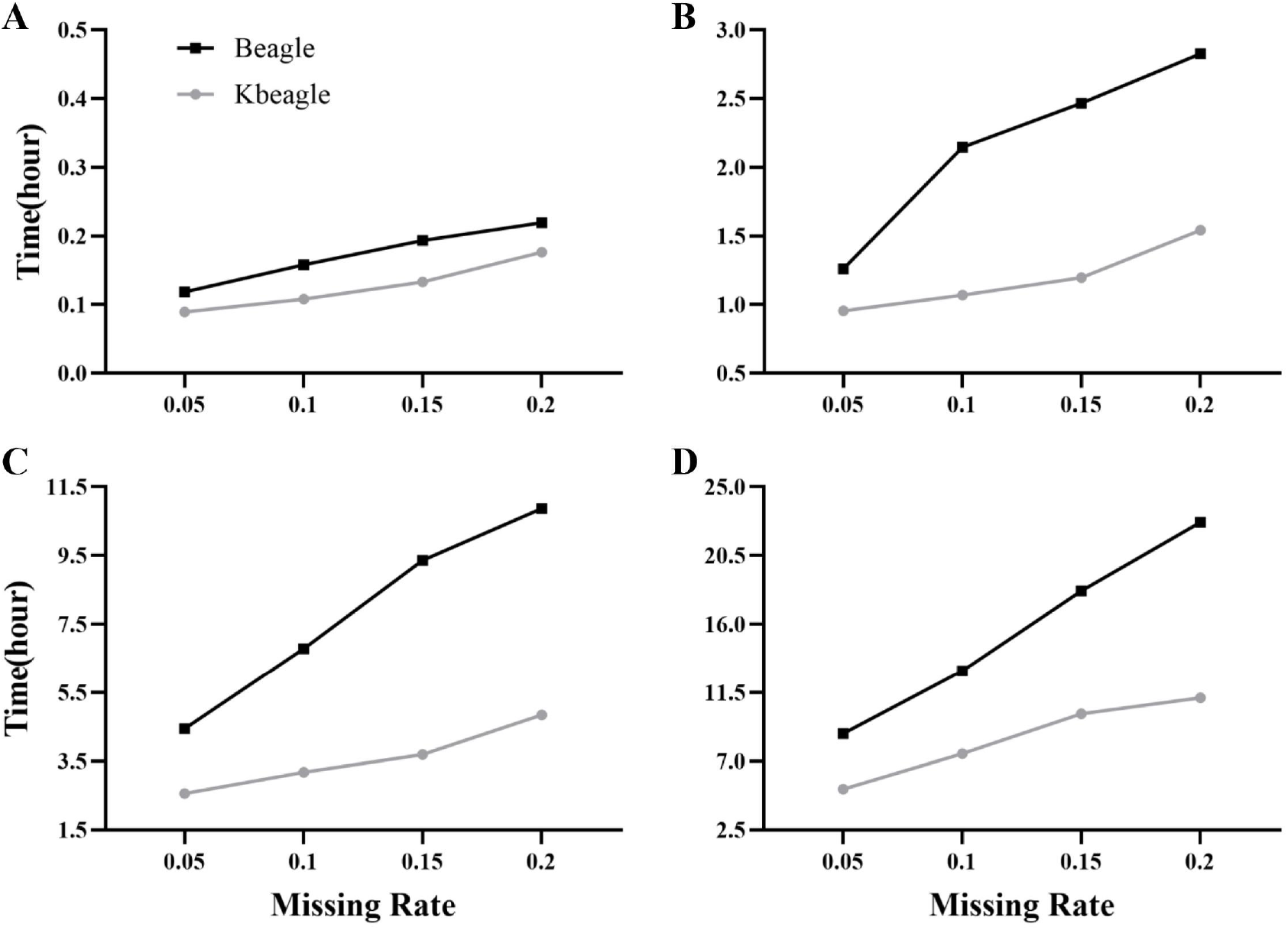
The computing time between Beagle software and KBeagle method. 10K(A), 30K(B), 50K(C) and 70K(D) data were randomly selected from the 98K data of 354 yaks. When using Beagle software and KBeagle method for genotype imputation, ‘system.time’ function was used to record the computing time of the two methods, including data inputting, data processing, genotype imputation and calculation of imputation accuracy. The function system.time started timing from the data inputting until the imputation accuracy was calculated, the total time of all steps was the computing time. The y-axis in each figure represented the imputation time (hour), the x-axis represented the missing rate, and the Beagle (blue broken line) and KBeagle (red broken line) were displayed as the average of 30 times replications. Obviously, the computing time of KBeagle method was less than that of Beagle software.

Under the same missing rate, considering the results at 10K (Fig. 3A), 30K (Fig. 3B), 50K (Fig. 3C), and 70K (Fig. 3D), the computing time of theKBeagle method was always less than that of the Beagle method. The computing time of the KBeagle and Beagle methods increased with the increase in the missing rate. The computing time of the Beagle method increased rapidly as the missing rate increased, while the computing time of the KBeagle method increased slower as the missing rate increased. The computing time of 10K data with a missing rate of 5% using the KBeagle method was the shortest at 0.09 hours. The longest computing time was 22.68 hours for 70K data with a missing rate of 20% using the Beagle method.

### Heritability

The heritability of the real data, the data containing the missing genotypes, and the imputed data were calculated using the GAPIT software package. Each dataset was repeated 30 times, and the average value was taken as the final heritability (Fig 4). The heritability of the real data was 0.801, whilst that of the data containing the missing genotype was 0.929, and that of the imputed data was 0.814. Therefore, this demonstrates that the heritability of the imputed data was close to that of the real data, while the heritability of the data containing the missing genotype was much higher than that of the other two datasets.

**Figure 4.**
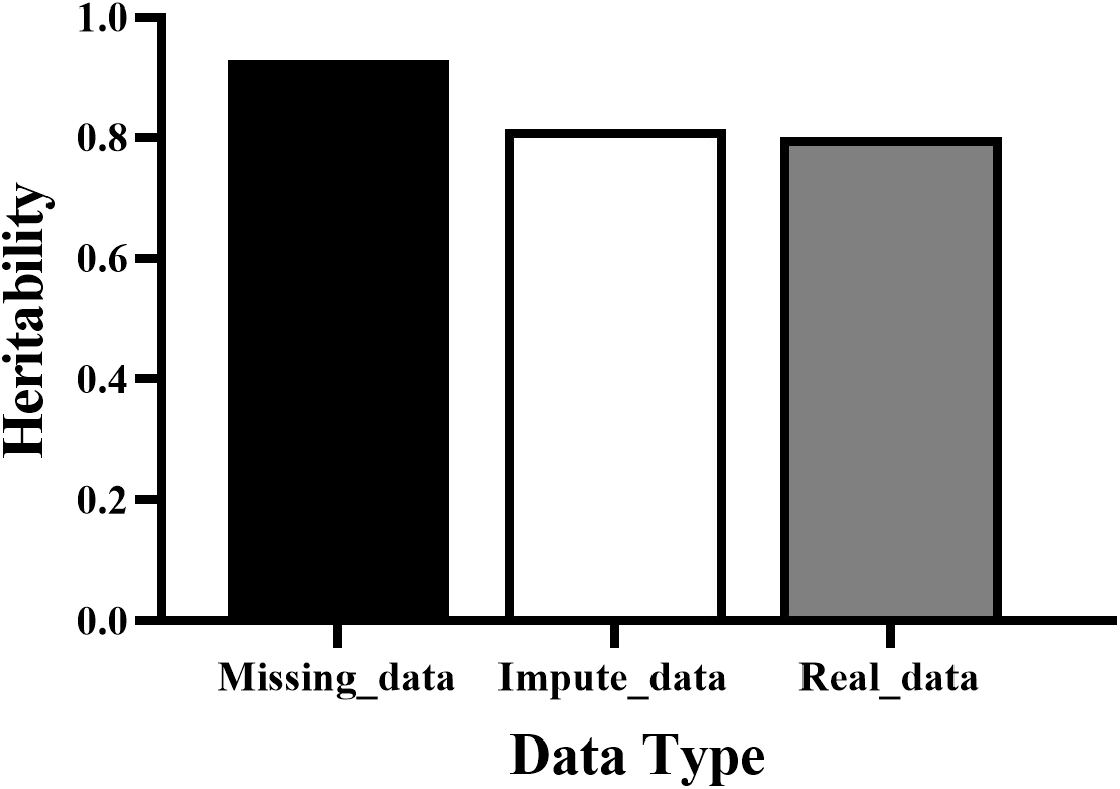
Comparison of estimated heritabilities with real, missing, and imputed genotypedata. Whole genotype data of 98K original data were used to simulate a trait for estimation of heritability. First, the missing rate of each SNPs was calculated, and the missing rate was ranked from high to low. Total 30 times replications were performed following simulation parameters. In the simulation, the heritability was set as 75%, thenumberofQTNwassetas20,andthetop20geneswereselectedfromSNPswith the highest missing rate.. The heritability was estimated using GAPIT software package. The ratio of estimated genetic variance between total phenotype variance was defined as estimated heritability.All compared datasets included three individual datasets: impute data (red bar) containing imputed genotype and others real genotypes, missing data (green bar) containing random imputed genotype (1, numeric genotype values means heterozygotes) and others real genotypes and original data (blue bar) containing whole real genotypes. The imputation method was KBeagle approach.

### Prediction accuracy

The mixed.solve (rrBLUP package in R) function was used to calculate the prediction accuracy of the real data, data containing missing genotypes, and the imputed data for genome selection analysis, and used the average value as the prediction accuracy (Fig S1). The prediction accuracy of the real data was 0.304, similar to that of the data containing the missing genotype at 0.303, and that of the imputed data at 0.305. The prediction accuracy of the three kinds of data was relatively low, with almost no difference between them.

## Discussion

Genotype imputation was performed using marker information from the linkage disequilibrium (LD) fragment. The estimated accuracy of fragments between individuals with known and unknown genotypes is the key factor in imputation ability. Yaks have been randomly mating with individuals in such populations or with wild yaks based on grazing on the plateau meadow. Therefore, the half LD decay of the yak population is much lower than that of other animals (Qiu et al. 2015; Wang et al. 2014; Bomba et al. 2015).In this study, the imputation accuracy of the Beagle software and the KBeagle method was approximately 85%–89%, which is lower than 90% (accuracy in other animals). Similar results were reported by Yang et al. (2020), where the imputation accuracy of Buffalo using the Beagle software was approximately 83.5%–84.5%. Yang used the 3300K reference panel of Buffalo to impute 1600K target samples, the imputation accuracy of the Beagle software was 84%, whilst the imputation accuracy of the Minimac3 software was 90%.

In this study, the genotype imputation method was used to impute the 10K, 30K, 50K, and 70K data of 354 Ashdan yaks, with the smaller reference panel having lower imputation accuracy than that of the larger reference plane. The study of Carvalheiro et al. (2014) showed that using 7K reference panels had lower imputation accuracy than when using 15K, 20K,and 75K reference panels. The reference panel in this study was small, and the imputation accuracy was relatively low, but as the reference panel increased, the imputation accuracy also gradually increased. Therefore, the results showed that the size of the reference panel could affect the imputation accuracy.

The data used in this study came from 354 Ashdan yaks in Datong Ranch. These samples were from a large group, and related to each other, and had family relationships. In a research report by Korkuć et al. (2019), the correlation between samples had a positive impact on the imputation accuracy from one German Black Pied cattle to another German Black Pied cattle. However, because the individuals in this study came from Datong Ranch, and mated randomly, their kinship was unclear, and the Beagle software could only assign genotypes to all, and could not cluster samples with high correlation. Therefore, the imputation accuracy was relatively poor. The KBeagle method clustered samples with higher correlation, and so, the samples within the same cluster had a closer genetic relationship and a better imputation effect. In the research report by Larmer et al. (2017), when using PLINK clustering software, the imputation accuracy of one or two imputation steps (directly from low density or from low to medium to high) was slightly higher than that of imputing directly using all samples. This showed the feasibility of clustering samples with high correlation and then imputing them.

In this study, the average imputation accuracy was taken as the final result, rather than the maximum or minimum imputation accuracy. The average imputation accuracy of the KBeagle method was 87%–89%, a little higher than the imputation accuracy of the Beagle software, but no significant effect was obtained. The KBeagle method used in this study could improve the imputation accuracy, but it needed population imputation with a large number of samples. Furthermore, the best cluster group calculated by the K-Means algorithm only found the cluster group that was best suited to all samples, but the average interpolation accuracy of these cluster groups was not necessarily the highest. The best cluster group may have many clusters with a small number of samples and low imputation accuracy, resulting in a low average imputation accuracy. The key to further optimizing the KBeagle method was to find the best cluster number K to maximize the average value of imputation accuracy.

In most studies, the imputation accuracy is calculated using correlation (Antolín et al. 2017; Ros-Freixedes et al. 2020; Pausch et al. 2017), which only describes the linear relationship between real and estimated alleles, rather than a measure of distance. Additionally, imputation accuracy can also be evaluated using other more complex methods such as calculating the rate of correctly imputing genotypes (percent identity), relative Manhattan distance (Zhang and Druet 2010), and calibration of the posterior genotype probability (Browning and Browning 2009). The relative Manhattan distance could explain different types of errors, and for some alleles with correct imputing (A/T imputing was A/A or T/T), the relative Manhattan distance thought that this should also be calculated in the imputation accuracy. Therefore, the imputation accuracy calculated by the distance from Manhattan was relatively high. In this study, percent identity was taken as the imputation accuracy. Since this method was relatively simple to calculate, and only identical genotypes could be considered as correct imputing the imputation accuracy calculated by percent identity was relatively low. However, the percent identity was closer to the real imputing situation, consistent with the results of most studies, making it more suitable for calculating the imputation accuracy.

Because there was no yak reference panel in the Beagle software, this study used the imputed population itself to impute the missing genotypes. When the Beagle software or the KBeagle method was used for imputing, all the samples except those samples to be imputed were reference panels. Both imputation methods used the Beagle software for imputation, and had the same parameters such as burning, iterations, and phase-states. The Beagle software used all samples as the reference panel, and it needed to infer the missing genotype based on the information from all samples, making the calculation process more complicated and time-consuming. However, after clustering, the KBeagle method had a smaller reference panel, used relatively less sample information when imputing, had a simpler calculation process, and took a long time. There was no simple linear relationship between the computing time and the size of the reference panel, so the time taken to divide all samples into multiple clusters for imputing was still less than the time taken to impute all samples directly. The KBeagle method could not only improve the imputation accuracy but also reduce the computing time, which was better than the Beagle software.

This study calculated the heritability of imputed data, data containing missing values, and real data. The heritability of imputed data and real data were higher than those for data containing missing values. When calculating the heritability of data containing missing values, the missing genotype was replaced with genotype 1, and so a large number of wrong genotypes led to high heritability. However, after imputing the imputed data, the wrong genotype was greatly reduced, and closer to the real data, which in turn resulted in the heritability being closer to that of the real data. This indicated that genotype imputation could affect the heritability of genome selection, and the imputed data could be used to better find the phenotypes with high heritability in species.

This study compared the prediction accuracy of genome selection with imputed data, data containing missing values, and real data. The results showed that the prediction accuracy of the three data types was relatively low and essentially the same, and that the prediction accuracy of imputed data was not significantly improved. In the research report by Weng et al.(2012), they showed that different imputation methods and quality control standards had no significant effect on the accuracy of genome selection, however, imputing before quality control could slightly improve the accuracy of genome selection. The number of samples in the reference panel and the number of SNPs of known genotypes in the genome could affect the prediction accuracy of genome selection (Hayes et al. 2009). The data used in this study were obtained after quality control, excluding SNPs with a call rate less than 90% and minor allele frequency (MAF) less than 1%, and individuals with a genotype rate less than 90% were also excluded. Therefore, the data was reduced after quality control, which may be the reason for the low accuracy of genome selection.

Compared with the Beagle software, the KBeagle method had higher imputation accuracy and shorter computing time. Next, genome selection was carried out for the data imputed by the KBeagle method, and its heritability and prediction accuracy were calculated to explore the imputation application of the KBeagle method. We found that the heritability was similar to that of the real data, and the prediction accuracy was relatively low. The results of this study showed that KBeagle method had a good imputation application, which provided a new direction for the study of genotype imputation.

## Acknowledgments

We would like to thank Editage (www.editage.cn) for English language editing. This project was partially funded by the Qinghai Science and Technology Program, China (2022-NK-110), Sichuan Science and Technology Program, China (Award #s 2021YJ0269 and 2021YJ0266), the Fundamental Research Funds for the Central Universities, China (Southwest Minzu University, ZYN2022023), and the Program of Chinese National Beef Cattle and Yak Industrial Technology System, China (CARS-37).

## Author Contributions

JQ: Software, Writing first draft of manuscript, Visualization, Testing, and Validation. XL: Data curation and Visualization. YL: Review and Editing. WP, YK, and JZ: Manuscript Revision. JW: Methodology, Supervision, Conceptualization and Manuscript Revision. All authors have read and approved the final manuscript.

## Competing Interests

The authors declare that they have no competing interests.

## Notes

### Competing Interest Statement

The authors have declared no competing interest.

## Reference

Antolín R, Nettelblad C, Gorjanc G, Money D and Hickey JM (2017) A hybrid method for the imputation of genomic data in livestock populations. Genet Sel Evol 49:1–17

Bomba L, Nicolazzi EL, Milanesi M, Negrini R, Mancini G, Biscarini F et al. (2015) Relative extended haplotype homozygosity signals across breeds reveal dairy and beef specific signatures of selection. Genet Sel Evol 47:1–14

Browning BL and Browning SR (2009) A unified approach to genotype imputation and haplotype-phase inference for large data sets of trios and unrelated individuals. Am J Hum Genet 84:210–223

Browning BL, Zhou Y and Browning SR (2018) A One-Penny Imputed Genome from Next-Generation Reference Panels. Am J Hum Genet 103:338–348

Browning SR and Browning BL (2007) Rapid and accurate haplotype phasing and missing-data inference for whole-genome association studies by use of localized haplotype clustering. Am J Hum Genet 81:1084–1097

Carvalheiro R, Boison SA, Neves HH, Sargolzaei M, Schenkel FS, Utsunomiya YT et al. (2014) Accuracy of genotype imputation in Nelore cattle. Genet Sel Evol 46:1–11

Chen J and Shi X (2019) Sparse Convolutional Denoising Autoencoders for Genotype Imputation. Genes (Basel) 10:652

Chud TC, Ventura RV, Schenkel FS, Carvalheiro R, Buzanskas ME, Rosa JO et al. (2015) Strategies for genotype imputation in composite beef cattle. BMC Genet 16:1–10

Endelman JB (2011) Ridge regression and other kernels for genomic selection with R package rrBLUP. Plant Genome 4:250–255

Hayes BJ, Bowman PJ, Chamberlain AJ and Goddard ME (2009) Invited review: Genomic selection in dairy cattle: progress and challenges. J Dairy Sci 92:433–443

Howie BN, Donnelly P and Marchini J (2009) A flexible and accurate genotype imputation method for the next generation of genome-wide association studies. PLoS Genet 5:e1000529

Jia C, Li C, Fu D, Chu M, Zan L, Wang H et al. (2020) Identification of genetic loci associated with growth traits at weaning in yak through a genome-wide association study. Anim Genet 51:300–305

Korkuć P, Arends D and Brockmann GA (2019) Finding the Optimal Imputation Strategy for Small Cattle Populations. Front Genet 10:52

Larmer SG, Sargolzaei M, Brito LF, Ventura RV and Schenkel FS (2017) Novel methods for genotype imputation to whole-genome sequence and a simple linear model to predict imputation accuracy. BMC Genet 18:1–12

Li N and Stephens M (2003) Modeling linkage disequilibrium and identifying recombination hotspots using single-nucleotide polymorphism data. Genetics 165:2213–2233

Li Y. (2006) Mach 1.0: rapid haplotype reconstruction and missing genotype inference. Am J Hum Genet 79: 2290

Liu B, Zhang T, Li Y, Liu Z and Zhang Z (2021) Kernel Probabilistic K-Means Clustering. Sensors (Basel) 21:1892

Medina CA, Kaur H, Ray I and Yu LX (2021) Strategies to Increase Prediction Accuracy in Genomic Selection of Complex Traits in Alfalfa (Medicago sativa L.). Cells 10:3372

Pausch H, MacLeod IM, Fries R, Emmerling R, Bowman PJ, Daetwyler HD et al. (2017) Evaluation of the accuracy of imputed sequence variant genotypes and their utility for causal variant detection in cattle. Genet Sel Evol 49:1–14

Purcell S, Neale B, Todd-Brown K, Thomas L, Ferreira MA, Bender D et al. (2007) PLINK: a tool set for whole-genome association and population-based linkage analyses. Am J Hum Genet 81:559–575

Qiu Q, Wang L, Wang K, Yang Y, Ma T, Wang Z et al. (2015) Yak whole-genome resequencing reveals domestication signatures and prehistoric population expansions. Nat Commun 6:1–7

Ros-Freixedes R, Whalen A, Gorjanc G, Mileham AJ and Hickey JM (2020) Evaluation of sequencing strategies for whole-genome imputation with hybrid peeling. Genet Sel Evol 52:1–19

Rubinacci S, Delaneau O and Marchini J (2020) Genotype imputation using the Positional Burrows Wheeler Transform. PLoS Genet 16:e1009049

Scheet P and Stephens M (2006) A fast and flexible statistical model for large-scale population genotype data: applications to inferring missing genotypes and haplotypic phase. Am J Hum Genet 78:629–644

Song H, Ye S, Jiang Y, Zhang Z, Zhang Q and Ding X (2019) Using imputation-based whole-genome sequencing data to improve the accuracy of genomic prediction for combined populations in pigs. Genet Sel Evol 51:1–13

Stachowicz K, Larmer S, Jamrozik J, Moore SS, Miller SP (2013) Sequencing and genotyping for the whole genome selection in Canadian beef populations. Armidale: Association for the Advancement of Animal Breeding and Genetics 20:344–347

Uemoto Y, Sasaki S, Sugimoto Y and Watanabe T (2015) Accuracy of high-density genotype imputation in Japanese Black cattle. Anim Genet 46:388–394

Wang J and Zhang Z (2021) GAPIT Version 3: Boosting Power and Accuracy for Genomic Association and Prediction. Genomics Proteomics Bioinformatics 19:629–640

Wang K, Hu Q, Ma H, Wang L, Yang Y, Luo W et al. (2014) Genome-wide variation within and between wild and domestic yak. Mol Ecol Resour 14:794–801

Wang Q, Yu Y, Yuan J, Zhang X, Huang H, Li F et al. (2017) Effects of marker density and population structure on the genomic prediction accuracy for growth trait in Pacific white shrimp Litopenaeus vannamei. BMC Genet 18:1–9

Weng Z, Zhang Z, Ding X, Fu W, Ma P, Wang C et al. (2012) Application of imputation methods to genomic selection in Chinese Holstein cattle. J Anim Sci Biotechnol 3:1–57

Yang W, Yang Y, Zhao C, Yang K, Wang D, Yang J et al. (2020) Animal-ImputeDB: a comprehensive database with multiple animal reference panels for genotype imputation. Nucleic Acids Res 48:D659–D667

Ye S, Yuan X, Huang S, Zhang H, Chen Z, Li J et al. (2019) Comparison of genotype imputation strategies using a combined reference panel for chicken population. Animal 13:1119–1126

Zhang Z and Druet T (2010) Marker imputation with low-density marker panels in Dutch Holstein cattle. J Dairy Sci 93:5487–5494

