## Supplementary material for "KBeagle: An Adaptive Strategy and Tool for Improvement of Imputation Accuracy and Computing Efficiency": Table 1, Table 2

**Table 1: Imputation matching rate of 10K, 30K, 50K and 70K data at 5%, 10%, 15% and 20% missing rate.**

| Data | Method | Missing Rate | | | |
|---|---|---|---|---|---|
| | | **5%** | **10%** | **15%** | **20%** |
| 10K | Beagle | 86.26% | 86.16% | 86.07% | 85.98% |
| | KBeagle | 87.22% | 87.18% | 87.03% | 86.88% |
| 30K | Beagle | 87.48% | 87.21% | 86.93% | 86.68% |
| | KBeagle | 88.43% | 88.10% | 87.85% | 87.59% |
| 50K | Beagle | 88.05% | 87.74% | 87.36% | 86.98% |
| | KBeagle | 88.62% | 88.32% | 88.08% | 87.79% |
| 70K | Beagle | 88.37% | 88.27% | 87.66% | 87.26% |
| | KBeagle | 88.72% | 88.46% | 88.22% | 87.93% |

**Table 2: Imputation time (hours) of 10K, 30K, 50K and 70K data at 5%, 10%, 15% and 20% missing rate.**

| Data | Method | Missing rate | | | |
|---|---|---|---|---|---|
| | | **5%** | **10%** | **15%** | **20%** |
| 10K | Beagle | 0.12 | 0.16 | 0.19 | 0.22 |
| | KBeagle | 0.09 | 0.11 | 0.13 | 0.18 |
| 30K | Beagle | 1.26 | 2.15 | 2.47 | 2.83 |
| | KBeagle | 0.95 | 1.07 | 1.20 | 1.54 |
| 50K | Beagle | 4.45 | 6.79 | 9.37 | 10.87 |
| | KBeagle | 2.56 | 3.18 | 3.70 | 4.85 |
| 70K | Beagle | 8.81 | 12.93 | 18.18 | 22.68 |
| | KBeagle | 5.17 | 7.49 | 10.10 | 11.15 |
