## Supplementary material for "KBeagle: An Adaptive Strategy and Tool for Improvement of Imputation Accuracy and Computing Efficiency": Fig S1

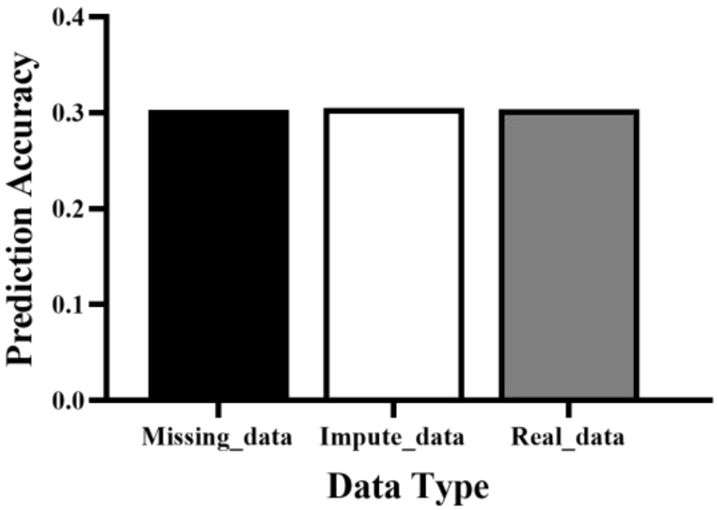


**Figure S1. The prediction ability of the genome selection of three data sets.**

The rrBLUP model was used to calculate the prediction ability of the imputed data , the compared data contains missing values, and real genotype data sets. The correlation between prediction phenotype values and simulated phenotype values in the inference population was used to define the prediction accuracy. The y-axis in the figure indicated the prediction ability, the x-axis indicated the data type, and the imputed data (red bar) and the data containing missing values (green bar) were displayed as the average value of 30 times replications. Each replicate used a five-fold cross-validation.
